# Measuring stimulus-evoked neurophysiological differentiation in distinct populations of neurons in mouse visual cortex

**DOI:** 10.1101/2020.11.27.400457

**Authors:** William G. P. Mayner, William Marshall, Yazan N. Billeh, Saurabh R. Gandhi, Shiella Caldejon, Andrew Cho, Fiona Griffin, Nicole Hancock, Sophie Lambert, Eric Lee, Jennifer Luviano, Kyla Mace, Chelsea Nayan, Thuyanh Nguyan, Kat North, Sam Seid, Ali Williford, Chiara Cirelli, Peter Groblewski, Jerome Lecoq, Giulio Tononi, Christof Koch, Anton Arkhipov

## Abstract

Despite significant progress in understanding neural coding, it remains unclear how the coordinated activity of large populations of neurons relates to what an observer actually perceives. Since neurophysiological differences must underlie differences among percepts, *differentiation analysis*—quantifying distinct patterns of neurophysiological activity—is an “inside out” approach that addresses this question. We used two-photon calcium imaging in mice to systematically survey stimulus-evoked neurophysiological differentiation in excitatory populations across 3 cortical layers (L2/3, L4, and L5) in each of 5 visual cortical areas (primary, lateral, anterolateral, posteromedial, and anteromedial) in response to naturalistic and phase-scrambled movie stimuli. We find that unscrambled stimuli evoke greater neurophysiological differentiation than scrambled stimuli specifically in L2/3 of the anterolateral and anteromedial areas, and that this effect is modulated by arousal state and locomotion. Contrariwise, decoding performance was far above chance and did not vary substantially across areas and layers. Differentiation also differed within the unscrambled stimulus set, suggesting that differentiation analysis may be used to probe the ethological relevance of individual stimuli.

## 1 Introduction

The visual system acts on incoming stimuli to extract meaningful features and guide behavior, a process that transforms physical input into conscious visual percepts. Since the early experiments of Hubel and Wiesel (1959), neuroscience has yielded considerable insight into the visual system by analyzing neural response properties to uncover which features cells are tuned to and how their activity relates to behavior. Modern decoding approaches have revealed stimulus information present in population responses (Quiroga & Panzeri, 2009). However, that a given population of neurons represents or encodes stimulus information does not imply that this information is used to generate conscious percepts in the subject (Brette, 2019). Consequently, despite the success of these “outside in” methods (Buzsáki, 2019) in understanding neural coding, it remains unclear how the coordinated activity of large populations of neurons relates to what the observer actually sees.

Is there an objective and quantitative approach to analyzing neural responses that can shed light on this question? *Differentiation analysis—*measuring the extent to which a population of neurons expresses a rich and varied repertoire of states—has been proposed as one such approach (Boly et al., 2015; Mensen et al., 2017, 2018). Differentiation analysis exemplifies “inside out” methodology (Buzsáki, 2019) in that the spatiotemporal diversity of neural activity (*neurophysiological differentiation* or ND) is quantified without reference to the stimulus or other experimental variables imposed *a priori* by the investigator, in contrast to feature tuning or decoding analyses. Supporting this proposal, recent studies in humans have shown that the ND evoked by a stimulus is correlated with subjective reports of its “meaningfulness” and the “number of distinct experiences” it elicits (Mensen et al., 2017, 2018).

A visual stimulus can be considered meaningful to the observer if it evokes rich and varied perceptual experiences (*phenomenological differentiation*). For example, an engaging movie is meaningful in this sense, as it evokes many distinct percepts with high-level structure; conversely, flickering ‘TV noise’ essentially evokes a single percept with no high-level structure to a human observer, even though, at the level of pixels, any two frames of noise are likely to be more different from each other than a pair of frames from a movie (*stimulus differentiation*). Since conscious percepts are generated by brain states, ND must underlie phenomenological differentiation. Thus one can expect to see correlations between ND and subjective perception of the “richness” or “meaningfulness” of stimuli, as has indeed been shown in human studies using fMRI and EEG (Boly et al., 2015; Mensen et al., 2017, 2018).

Moreover, integrated information theory (IIT) posits a fundamental relationship between ND and subjective experience itself. This theoretical framework predicts that consciousness requires the joint presence of integration and differentiation: that is, a system is conscious if it is causally irreducible to its parts and possesses a rich dynamical repertoire of states (Tononi, 2004; Oizumi et al., 2014; Tononi et al., 2016). Theoretical work has demonstrated that ND can serve as a proxy for integrated information in highly recurrent systems where integration can be assumed, such as the brain (Marshall et al., 2016). Consistent with IIT’s predictions, several studies have employed differentiation analysis across a range of species and spatiotemporal scales to show that loss of ND is implicated in loss of consciousness (Casali et al., 2013; Hudetz et al., 2014; Barttfeld et al., 2015; Wenzel et al., 2019).

However, although the applications of differentiation analysis cited above suggest that ND can provide a readout of stimulus-evoked phenomenological differentiation (Boly et al., 2015; Mensen et al., 2017, 2018), the low spatial resolution of fMRI and EEG has so far precluded identifying the specific cell populations that underlie this correspondence. Indeed, a longstanding question of fundamental importance is which populations of neurons contribute directly to generating conscious percepts (Koch et al., 2016; Tononi et al., 2016; Mashour et al., 2020). According to the considerations above, differentiation analysis can shed light on this question, but to do so it must be applied to recorded brain signals from specific populations of neurons. This requires systematically measuring stimulus-evoked ND with cellular resolution.

To address this gap, we leveraged the Allen Institute for Brain Science (AIBS) pipeline for *in vivo* two-photon calcium (Ca^2+^) imaging (de Vries et al., 2020) to measure stimulus-evoked ND in the visual cortex of the mouse. The present work represents one of the first projects within the *OpenScope* initiative, a collaborative model in which the capabilities of the AIBS “brain observatory” are made available to the wider neuroscientific community. The standardized, high-throughput OpenScope data acquisition pipeline allowed us to conduct a systematic survey of ND in excitatory cell populations across 3 cortical layers—layer (L) 2/3, L4, and L5—in each of 5 visual cortical areas: primary (V1), lateral (L), anterolateral (AL), posteromedial (PM), and anteromedial (AM), as awake mice were presented with visual stimuli. We used twelve 30 s movie stimuli chosen to span different levels of putative ethological relevance. The stimuli included naturalistic video clips of predators, prey, conspecifics, the home cage, movement through the underbrush of a forest (putatively of high ecological relevance to mice); clips of roadways, automobiles, and humans (putatively of low ethological relevance); and artificially generated clips with no ethological relevance. Some of the artificial stimuli were phase-scrambled versions of the naturalistic stimuli, which enabled us to contrast stimuli containing high-level structure against meaningless stimuli while controlling for low-order statistics.

We hypothesized that unscrambled naturalistic stimuli, which presumably elicit meaningful visual percepts, would evoke greater ND than their meaningless phase-scrambled counterparts. Indeed, we find that unscrambled stimuli evoke greater ND than scrambled stimuli specifically in L2/3 of areas AL & AM (*i.e.*, not in L4 or L5 of any area, nor in any sampled layer of areas V1, L, and PM). This effect is modulated by arousal and behavioral state and is robust to different methods of measuring ND. We contrast this layer- and area-specific finding with a decoding analysis that shows that information about the stimulus category, whether meaningful or meaningless, is present in most cell populations. This highlights a key difference between the methodological approaches: ND is more plausibly correlated with stimulus meaningfulness than the information measured by decoding, since the latter may not be functionally relevant (Brette, 2019). Furthermore, we find differences in evoked ND among the unscrambled stimuli that suggest that differentiation analysis can probe meaningfulness of individual stimuli.

## 2 Results

Using the AIBS OpenScope two-photon Ca^2+^ imaging pipeline (de Vries et al., 2020; **Figure 1A–D**), we recorded from the left visual cortex of mice while they passively viewed stimuli presented to the contralateral eye. We used the transgenic lines Cux2, Rorb, and Rbp4 (3 mice each) in which GCaMP6f is expressed in excitatory neurons predominantly in L2/3, L4, and L5, respectively. Visual cortical areas were delineated via intrinsic signal imaging (ISI; **Figure 1B**). Data were collected from 5 areas (V1, L, AL, PM, and AM**; Figure 1E**) across 45 experimental sessions (15 sessions per Cre line; 5 sessions per area; ~5 sessions per mouse). Mice were head-fixed and free to move on a rotating disc. Pupil diameter and running velocity were recorded. During each 70-minute session, twelve 30 s movie stimuli were presented in a randomized block design with 10 repetitions, with 4 s of mean-luminance grey shown between stimulus presentations (**Figure 1F,G**; see **Stimuli**). Stimuli were presented in greyscale but were not otherwise modified (in particular, it should be noted that spatial frequencies beyond the mouse acuity limit will appear blurred to the mice). Representative ΔF/F_0_ traces and behavioral data are shown in **Figure 1H**. One imaging session in L5 of AL was excluded from our analyses because of technical problems with the two-photon recording.

**Figure 1.**
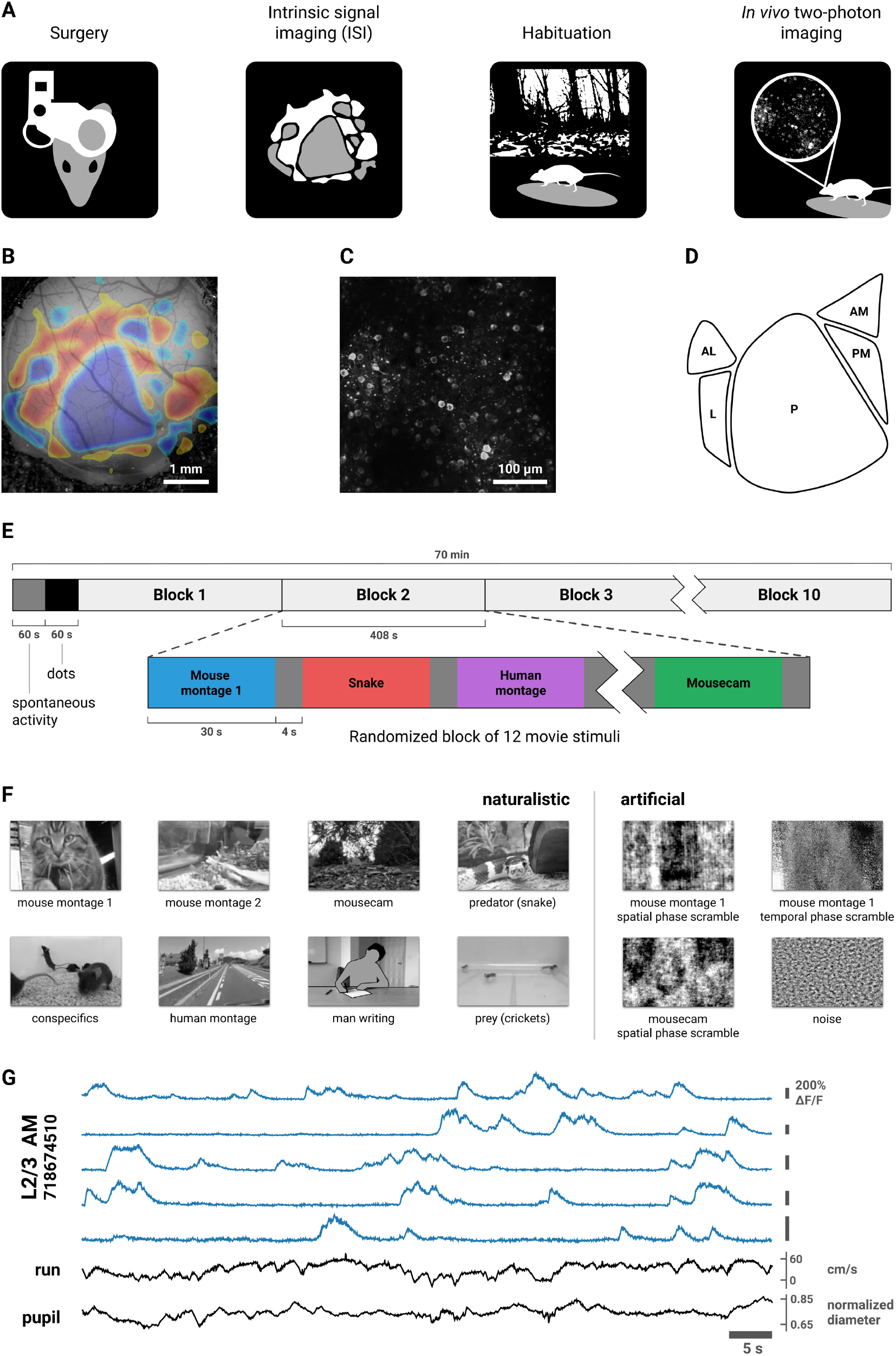
Experimental design. **(A)** Data were acquired using the AIBS’ standardized two-photon calcium imaging pipeline (de Vries et al., 2020; Groblewski et al., 2020; see **Methods**). Briefly, a custom headframe was implanted; intrinsic signal imaging (ISI) was performed to delineate retinotopically mapped visual areas; the mouse was habituated to the passive viewing paradigm over the course of ~2 weeks; and two-photon calcium imaging was performed in the left visual cortex while animals viewed stimuli presented to the contralateral eye in several experimental sessions. During the imaging sessions, head-fixed mice were free to run on a rotating disc. Locomotion velocity was recorded and pupil diameter was extracted from video of the animal’s right eye. **(B)** Example of an ISI map. **(C)** Example frame from a two-photon movie. Imaging data was processed as described in de Vries et al. (2020) to obtain ΔF/F_0_ traces. **(D)** Schematic of the 5 visual areas targeted in this study. **(E)** 10 randomized blocks of twelve 30 s movie stimuli were presented. 4 s of mean-luminance grey was presented between stimuli. The first 60 s was mean-luminance grey (spontaneous activity); the second 60 s period was a high-contrast sparse noise stimulus (not analyzed in this work). **(F)** Still frames from the 8 naturalistic (left) and 4 artificial (right) movie stimuli (see **Stimuli**). Two of the naturalistic stimuli, “mouse montage 1” and “mousecam”, were phase-scrambled to destroy high-level image features while closely matching low-order statistics (see **Phase scrambling**). **(G)** ΔF/F_0_ traces from 5 example cells, locomotion velocity, and normalized pupil diameter from a representative experimental session. *Note:* the “man writing” stimulus frame in (F) has been de-identified for presentation in this preprint in accordance with bioRxiv policy.

To measure ND, we employed a method from Mensen et al. (2018) for analyzing a set of timeseries recorded during the presentation of a continuous stimulus (**Figure 2**). Briefly, the power spectrum of each cell’s ΔF/F_0_ trace was estimated in 1 s windows. The cells’ power spectra during simultaneous windows were concatenated to form a vector representing the neurophysiological state of the population during that window. We calculated ND for each trial as the median Euclidean distance between the 30 population states elicited over the course of the 30 s stimulus. We computed distances in the frequency domain rather than the time domain in order to focus on differences in overall population state rather than differences in precise timing of ΔF/F_0_ transients. To account for variability in the size of the imaged populations we divided ND values by the square root of the number of cells (see **Spectral differentiation**). Spectral differentiation will be zero when the set of ΔF/F_0_ traces is perfectly periodic with a period of 1 s (the window size), and it will be high when many traces exhibit temporally varied patterns across the 30 seconds. The measure scales with the magnitude of the signal and thus has no well-defined maximum.

**Figure 2.**
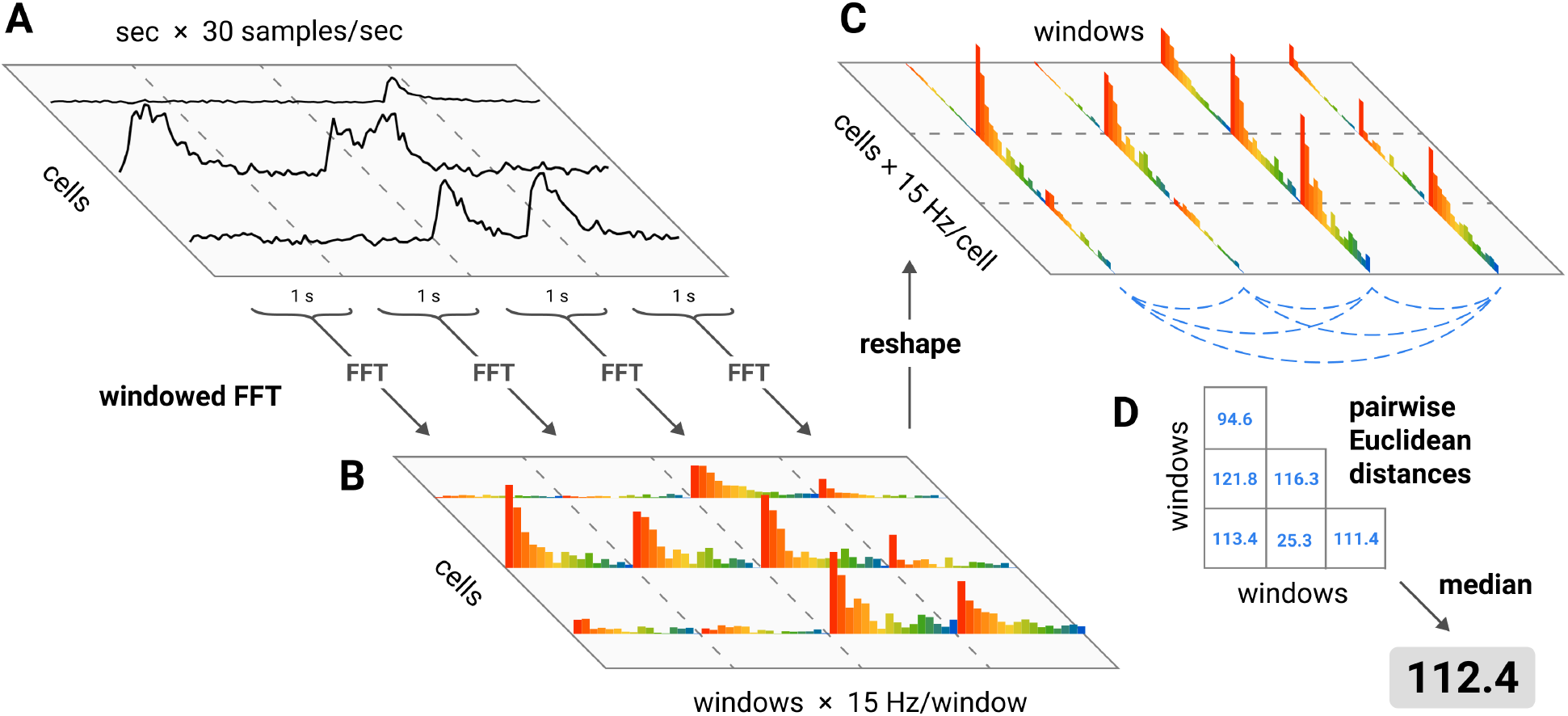
Spectral differentiation analysis. ND was computed as follows: **(A)** for each cell, the ΔF/F_0_ trace of each cell during stimulus presentation was divided into 1 s windows; **(B)** the power spectrum of each window was estimated; **(C)** the “neurophysiological state” during each 1 s window was defined as a vector in the high-dimensional space of cells and frequencies (*i.e.*, the concatenation of the power spectra in that window for each cell); **(D)** the ND in response to a given stimulus was calculated as the median of the pairwise Euclidean distances between every state that occurred during the stimulus presentation.

To compare the differentiation of responses to naturalistic and artificial stimuli, we generated Fourier phase-scrambled versions of two of our movie stimuli (**Figure 1G**). Phase-scrambling destroys the natural structure of the stimulus while closely matching the power spectrum (the spectrum was not conserved exactly because of numerical representational limitations of the stimulus format; see **Phase scrambling**). Note that operations that leave the power spectrum of a signal unchanged will not affect its spectral differentiation.

For the “mouse montage 1” stimulus (a montage of six 5 s naturalistic movie clips), we performed the phase-scrambling in two ways: (1) along the temporal dimension, on each pixel independently, and (2) along the two spatial dimensions, on all pixels. For the “mousecam” stimulus (a continuous 30 s clip of movement at ground level through the underbrush of a forest) we performed only the spatial phase-scrambling. This yielded 2 unscrambled stimuli and 3 scrambled stimuli. The full set of twelve stimuli was designed to span different levels of putative ethological relevance; here, we focus on the comparison of the unscrambled stimuli to their scrambled versions because low-order stimulus statistics are controlled and thus the contrast can be more easily interpreted.

### 2.1 Unscrambled stimuli elicit more differentiated responses compared to scrambled stimuli

We hypothesized that the unscrambled stimuli would elicit higher ND than their phase-scrambled counterparts. We tested this by fitting linear mixed effects (LME) models with experimental session as a random effect (see **Linear mixed effects models**); mean differences in ND of responses to unscrambled vs. scrambled stimuli are shown in **Figure 3**. We obtained similar results contrasting naturalistic vs. artificial stimuli across the entire stimulus set (**Supplementary Figure 1**). ND values were approximately log-normally distributed, so we applied a logarithmic transform to ND in all statistical analyses (see **Statistics**).

**Figure 3.**
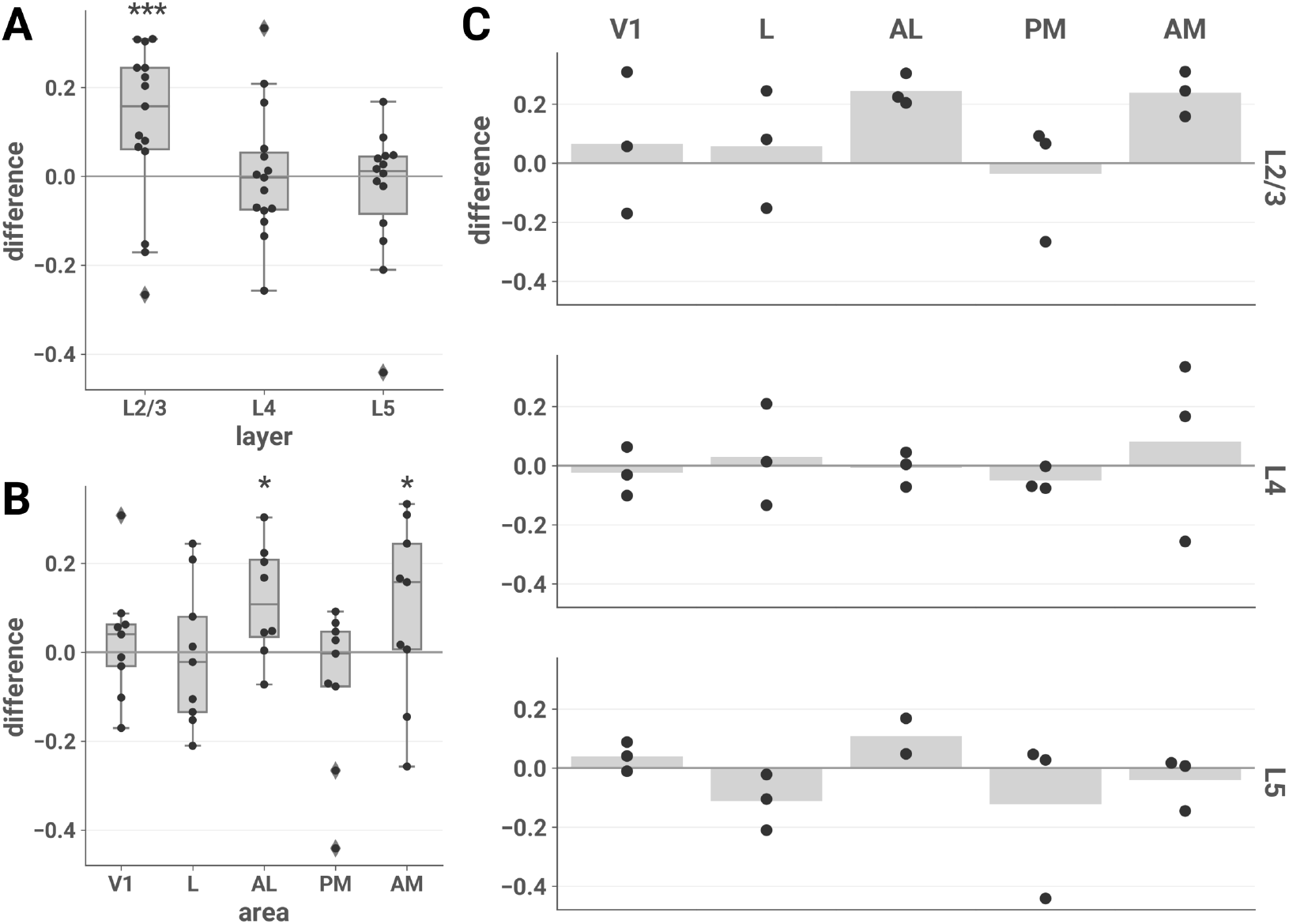
ND elicited by unscrambled vs. scrambled stimuli is higher in L2/3 of areas AL and AM. The mean difference in ND of responses to unscrambled vs. scrambled stimuli is plotted for each session by layer (A), area (B), and layer-area pair (C). **(A)** and **(B)**: asterisks indicate significant post hoc one-sided *z*-tests in the layer (A) and area (B) interaction LME models (*, p < 0.05; ***, p < 0.001). Boxes indicate quartiles; whiskers indicate the minimum and maximum of data lying within 1.5 times the inter-quartile range of the 25% or 75% quartiles; diamonds indicate observations outside this range. **(C)** Mean values are indicated by bars.

#### 2.1.1 Increased differentiation for unscrambled stimuli is specific to excitatory cells in L2/3

We found that unscrambled stimuli elicited more differentiated responses specifically in L2/3 (**Figure 3A**). We fitted an LME model with stimulus category (unscrambled or scrambled), layer, and their interaction as fixed effects and found a significant interaction (likelihood ratio test, *χ^2^*(2) = 13.379, p = 0.00124). Post hoc tests showed that the unscrambled vs. scrambled difference was specific to L2/3 (one-sided *z*-test; L2/3, z = 3.866, p = 1.66e–4; L4, z = 0.191, p = 0.810; L5, z = –1.168, p = 0.998; adjusted for multiple comparisons).

#### 2.1.2 Increased differentiation for unscrambled stimuli is specific to areas AL and AM

The increased ND in response to unscrambled stimuli was area-specific (**Figure 3B**). We fitted an LME model with stimulus category, area, and their interaction as fixed effects and found a significant interaction (likelihood ratio test, *χ^2^*(4) = 15.202, p = 0.00430). Post hoc tests showed that the unscrambled vs. scrambled difference was specific to AL and AM (one-sided *z*-test; V1, z = 0.704, p = 0.7479; L, z = –0.234, p = 0.9887; AL, z = 2.873, p = 0.0101; PM, z = –1.843, p > 0.999; AM, z = 2.446, p = 0.0356; adjusted for multiple comparisons).

### 2.2 Permutation tests for individual experimental sessions

The above analysis shows that the mean ND elicited by unscrambled stimuli is greater than for their phase-scrambled counterparts, and that this effect is driven by L2/3 cells in areas AL and AM. We also analyzed ND at the level of individual sessions with non-parametric permutation tests. For each session, we obtained a null distribution by randomly permuting the trial labels (unscrambled or scrambled) 20,000 times and computing the difference in mean ND on unscrambled vs. scrambled trials for each permutation. P values were computed as the fraction of permutations for which the permuted difference was greater than the observed difference.

The results of the individual session analyses were consistent with the LME analyses (**Table 1**). In all sessions recorded from L2/3 of AL & AM, responses to unscrambled stimuli were significantly more differentiated than to scrambled stimuli (p < 0.05).

**Table 1.** Permutation tests show increased ND for unscrambled vs. scrambled stimuli in L2/3 of AL & AM at the level of individual experimental sessions. Entries contain the fraction of sessions in which the mean ND of responses to unscrambled stimuli was significantly greater than responses to their scrambled counterparts vs. total number of sessions at a threshold of α = 0.05. For each session, a null distribution was obtained by randomly permuting trial labels (unscrambled or scrambled) 20,000 times and computing the difference in mean ND on unscrambled and scrambled trials for each permutation. P values were computed as the fraction of permutations for which the permuted difference was greater than the observed difference.

|  | V1 | L | AL | PM | AM | All areas |
| --- | --- | --- | --- | --- | --- | --- |
| **L2/3** | 1 / 3 | 1 / 3 | 3 / 3 | 0 / 3 | 3 / 3 | 8 / 15 |
| **L4** | 0 / 3 | 1 / 3 | 0 / 3 | 0 / 3 | 0 / 3 | 1 / 15 |
| **L5** | 0 / 3 | 0 / 3 | 0 / 2 | 0 / 3 | 0 / 3 | 0 / 14 |
| **All layers** | 1 / 9 | 2 / 9 | 3 / 8 | 0 / 9 | 3 / 9 |  |

Locomotion and pupil diameter can be considered behavioral indications of engagement with the environment (Jacobs et al., 2018; Ganea et al., 2018; Bennett et al., 2013) and modulate neuronal activity in visual cortex (Dadarlat & Stryker, 2017; McGinley, David, et al., 2015; McGinley, Vinck, et al., 2015; Niell & Stryker, 2010; Polack et al., 2013; Reimer et al., 2014; Salkoff et al., 2020; Vinck et al., 2015). We found that in L2/3 of AL & AM, effect sizes were positively correlated with locomotion activity (**Figure 4**, top left; Pearson’s *r* = 0.896; two-sided *t*-test; *t*(4) = 4.030, p = 0.0157) and pupil diameter (**Figure 4**, top right; *r* = 0.716; *t*(42) = 2.054, p = 0.109), suggesting that the difference in ND is more clear when the animal is engaged with the stimuli. This pattern was not evident when considering all cell populations (locomotion activity: **Figure 4,** bottom left, *r* = –0.034 (*t*(42) = –0.220, p = 0.827); pupil diameter: **Figure 4,** bottom right, *r* = 0.047 (*t*(42) = 0.302, p = 0.764).

**Figure 4.**
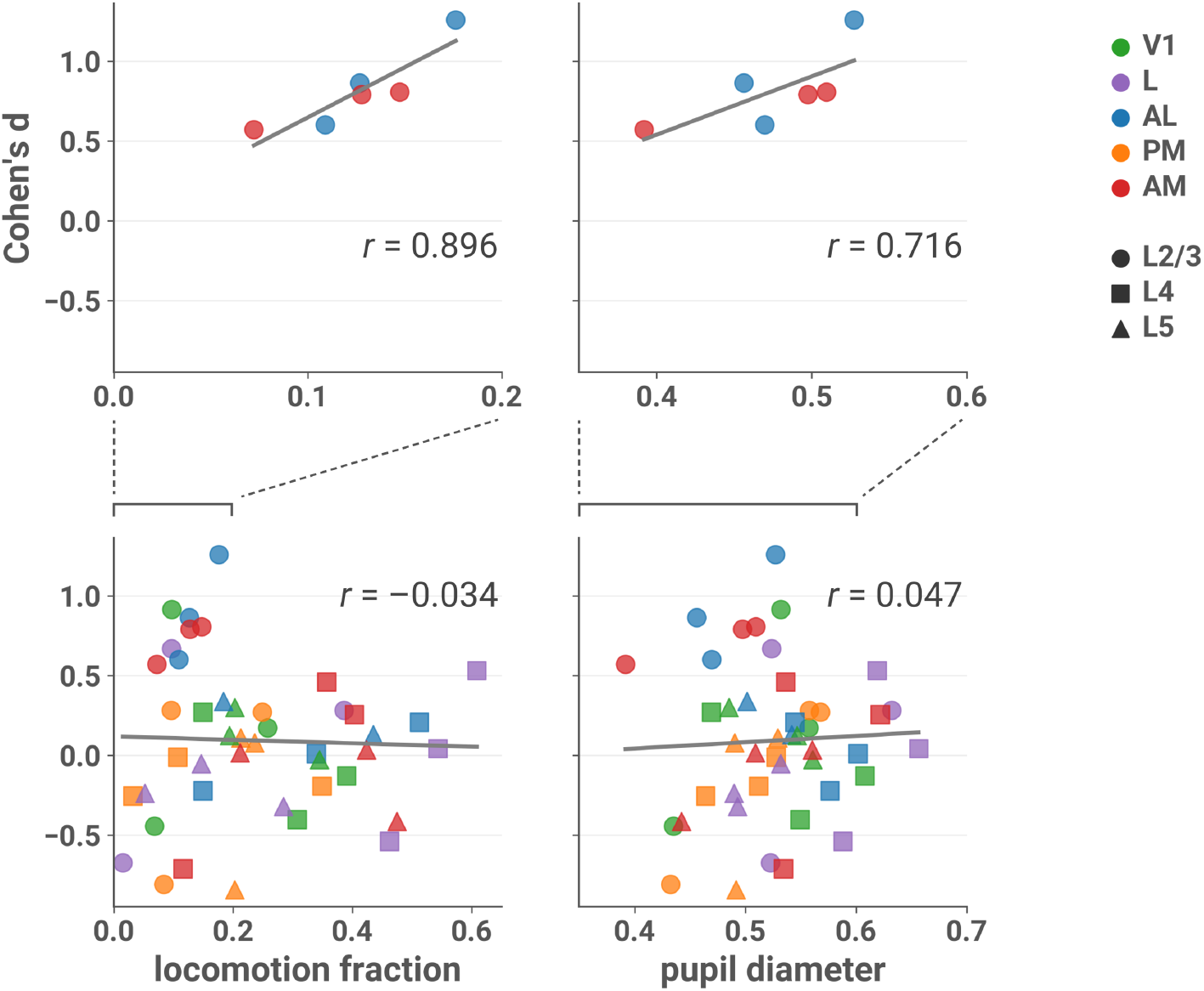
Effect sizes in L2/3 of AL & AM are larger in sessions with more locomotion and larger pupil diameter. Cohen’s *d* is plotted against the fraction of locomotion activity (left column) and mean normalized pupil diameter (right column) during the session, with linear fit in grey. Top row: only sessions recorded from L2/3 and areas AL or AM; bottom row: all sessions (note different scales). Top left, Pearson’s *r* = 0.896 (two-sided *t*-test; *t*(4) = 4.030, p = 0.0157); top right, *r* = 0.716 (*t*(42) = 2.054, p = 0.109); bottom left, *r* = –0.034 (*t*(42) = –0.220, p = 0.827); bottom right, *r* = 0.047 (*t*(42) = 0.302, p = 0.764). Running velocity greater than 2.5 cm/s was considered locomotion activity (see **Locomotion**). Normalized pupil diameter was obtained by dividing by the maximum diameter that occurred during the session (see **Pupillometry**).

### 2.3 Multivariate analysis also shows increased differentiation for unscrambled stimuli

Spectral differentiation is a univariate measure sensitive to differences within a given cell’s responses across time. To ensure that our results were not due to this particular measure, we also employed a multivariate approach that considers spatiotemporal differences in activity patterns across the cell population. For each session, the dimensionality of the population response vectors was reduced to 10 using UMAP (McInnes et al., 2018). In the resulting 10-dimensional space, ND was measured as the mean Euclidean distance to the centroid of the set of responses corresponding to that stimulus (see **Multivariate differentiation**).

The results of the multivariate analysis were consistent with those found using the spectral differentiation measure. The mean centroid distance was higher in response to unscrambled compared to scrambled stimuli (**Figure 5**), and this effect was specific to L2/3 (layer × stimulus category interaction: likelihood ratio test, *χ^2^*(2) = 8.263, p = 0.0161; post hoc one-sided *z*-tests: L2/3, z = 3.610, p = 0.000459; L4, z = 0.397, p = 0.720; L5, z = –0.202, p = 0.926) and areas AL and AM (area × stimulus category interaction: likelihood ratio test, *χ^2^*(4) = 15.659, p = 0.00351; post hoc tests: V1, z = –0.259, p = 0.990; L, z = –0.828, p > 0.999; AL, z = 2.546, p = 0.0270; PM, z = –0.051, p = 0.975; AM, z = 3.668, p = 0.000612).

**Figure 5.**
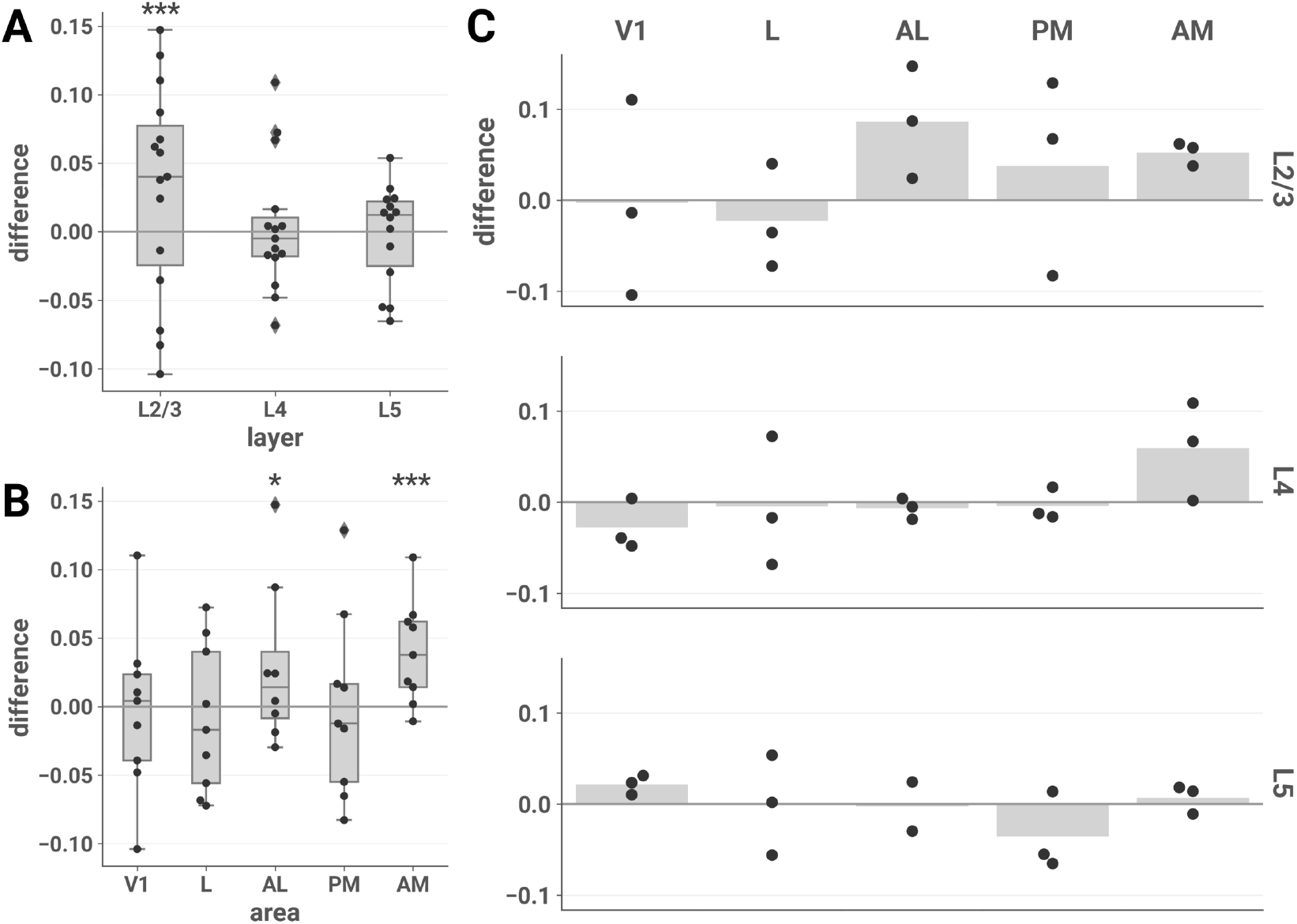
Multivariate differentiation analysis. The mean difference in the mean centroid distance of responses to unscrambled vs. scrambled stimuli is plotted for each session by layer (A), area (B), and layer-area pair (C). ND elicited by unscrambled vs. scrambled stimuli is higher in L2/3 and areas AL and AM, consistent with the spectral differentiation analysis. **(A)** and **(B)**: asterisks indicate significant post hoc one-sided *z*-tests in the layer (A) and area (B) interaction LME models (***, p < 0.001). Boxes indicate quartiles; whiskers indicate the minimum and maximum of data lying within 1.5 times the inter-quartile range of the 25% or 75% quartiles; diamonds indicate observations outside this range. **(C)** Mean values are indicated by bars.

### 2.4 Decoding analysis does not reveal layer or area specificity

We next asked whether the layer and area specificity of our ND results would be reflected in our ability to decode the stimulus category (unscrambled or scrambled) from population responses. We performed fivefold cross-validated linear discriminant analysis to decode stimulus category for each session and scored the classifier using balanced accuracy (see **Decoding analysis**). Decoding performance was high for most areas and layers (**Figure 6**), in contrast to the unscrambled-scrambled difference in ND. Performance was also high across layers and areas when we decoded stimulus identity, rather than category, using responses to all 12 stimuli (**Supplementary Figure 1**).

**Figure 6.**
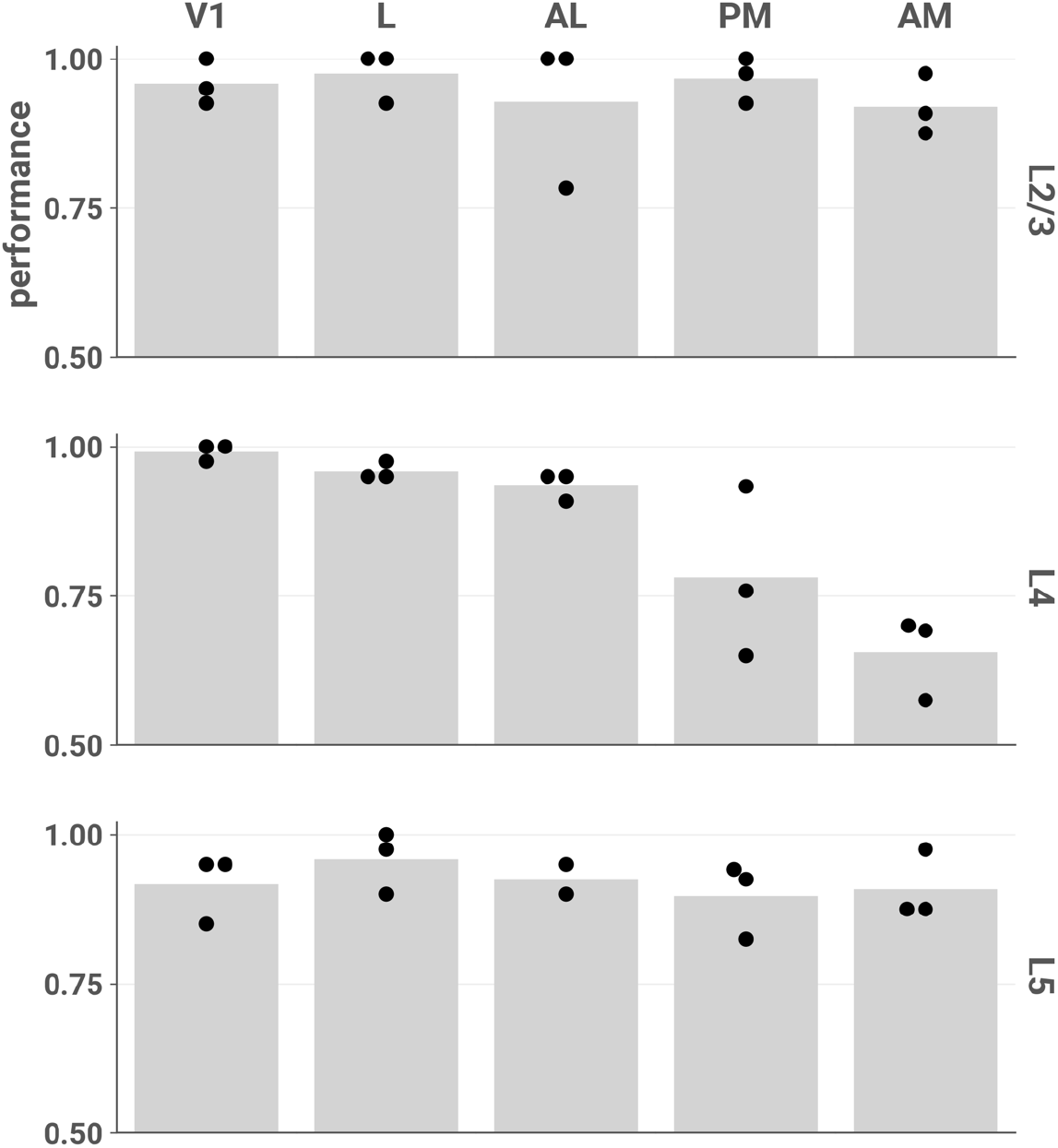
Stimulus category (unscrambled or scrambled) can be accurately decoded from most layers and areas. Each point represents the mean fivefold cross-validated balanced accuracy score of linear discriminant analysis performed on a single session (see **Decoding analysis)**. Chance performance is 0.5.

### 2.5 Differences in ND among individual stimuli

We also investigated whether ND differed among stimuli within the same category. This analysis was restricted to the set of unscrambled stimuli without jump cuts, *i.e.,* the 5 naturalistic continuous 30 s clips, to avoid potential confounds in comparing stimuli with and without abrupt transitions between different scenes. Here we used data from all layers and areas, since although L2/3 of AL & AM underlies unscrambled/scrambled differences, within-category differences might not be restricted to that subset. We fitted an LME model with stimulus as a fixed effect and found it was significant (likelihood ratio test, *χ^2^*(4) = 32.115, p = 1.812e–6). Post-hoc pairwise two-sided *t*-tests (adjusted for multiple comparisons), shown in **Figure 7**, revealed that the predator stimulus (a snake) evoked significantly higher differentiation than clips of conspecifics (*t*(2156) = 3.229, p = 0.0111); prey (crickets) (*t*(2156) = 3.928, p = 0.000839), and a man writing (*t*(2156) = 5.248, p = 1.670e–6). The “mousecam” clip of movement through a wooded environment also evoked a significantly higher differentiation than the clip of a man writing (*t*(2156) = 3.396 p = 0.00625). Here we present the main effect of stimulus; for an exploration of interactions with layer and area, and a comparison to decoding, see Supplementary Figure 3.

**Figure 7.**
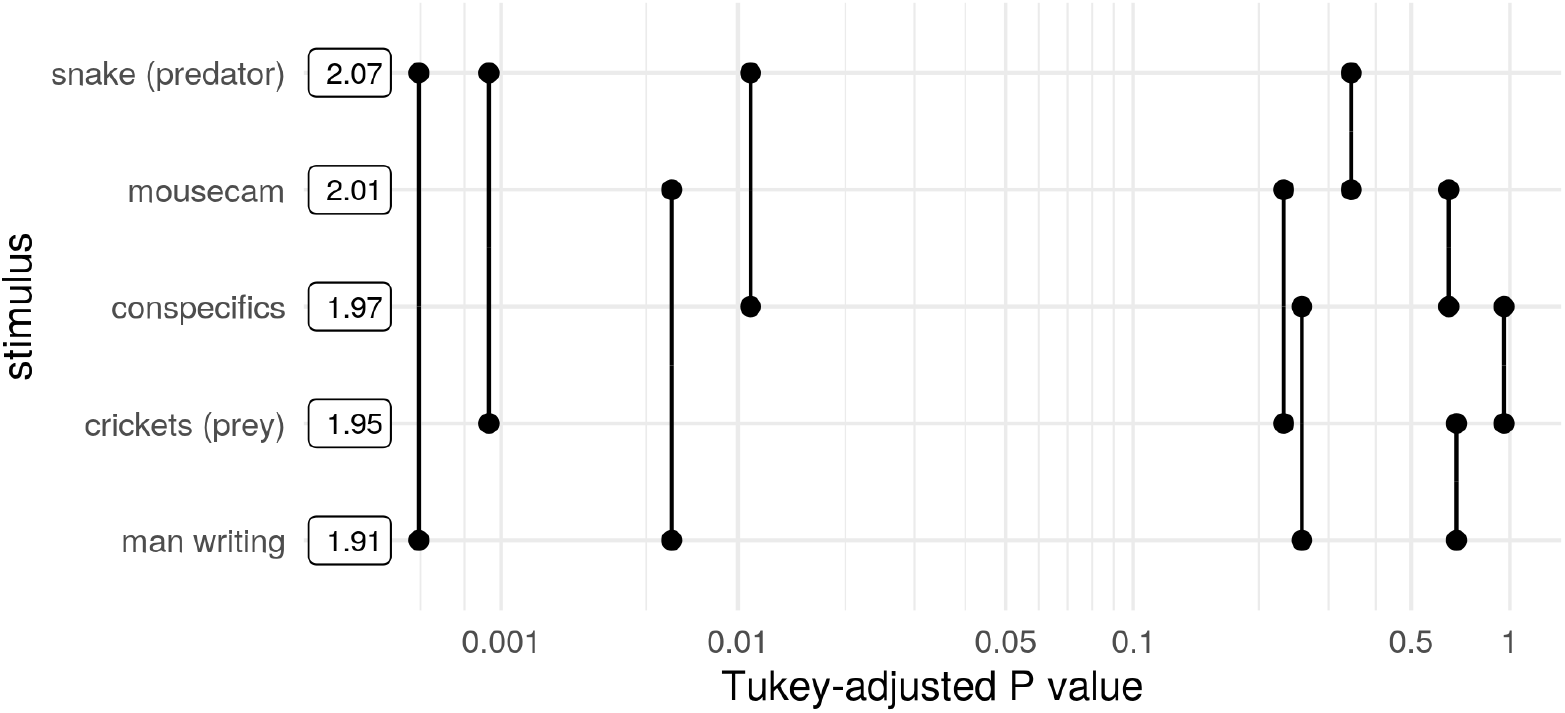
Pairwise differences in ND among unscrambled, continuous stimuli. Post hoc pairwise comparisons using data from all neuronal populations are plotted against their p values (adjusted for multiple comparisons). Boxes show mean ND for each stimulus. ND of the snake stimulus is significantly greater than that of crickets and man writing at a threshold of α = 0.01, and greater than conspecifics at α = 0.05. ND of the mousecam stimulus is greater than that of man writing at α = 0.01.

**Figure 8.**
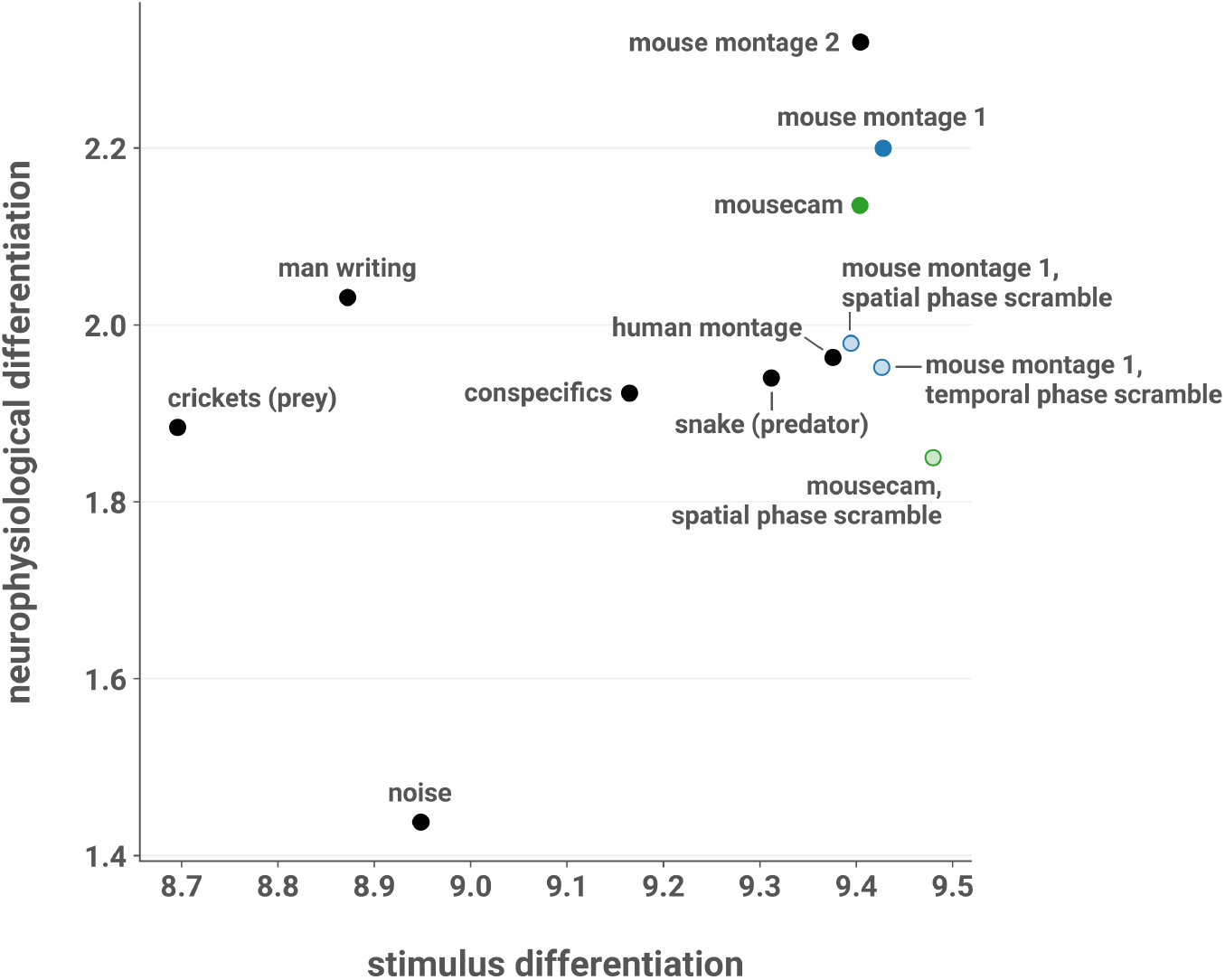
SD does not explain ND. Mean ND elicited by each stimulus in L2/3 of AL and AM, plotted against SD. SD was computed by treating each pixel of the movie as a “cell” and applying the spectral differentiation measure to traces of pixel intensities over time. Across all stimuli, mean ND is positively correlated with SD (Pearson’s *r* = 0.446; one-sided *t-*test; *t*(10) = 1.574, p = 0.0733). However, here the noise stimulus is an influential observation (Cook’s *D* = 0.665, more than twice as large as the next most influential observation). With the noise stimulus excluded, the correlation is weaker (*r =* 0.290; one-sided *t*-test; t(9) = 0.908, p = 0.194). Moreover, there was no evidence of a relationship with ND when considering only the scrambled stimuli and their unscrambled counterparts (*r* = –0.537; two-sided *t-*test; *t*(3) = –1.104. p *=* 0.350).

### 2.6 Stimulus differentiation does not explain ND

It is possible that ND does not reflect functionally relevant visual processing but is instead merely inherited from the differentiation of the stimulus itself. To rule out this possibility, we computed the stimulus differentiation (SD) by treating each pixel of the stimulus as a “cell” and applying the spectral differentiation measure to the traces of pixel intensities over time. Within L2/3 of AL and AM, the mean ND elicited by each stimulus was positively correlated with SD (Pearson’s *r* = 0.446, one-sided *t-*test; *t*(10) = 1.574, p = 0.0733). However, the noise stimulus is an influential observation (Cook’s *D* = 0.665, more than twice as large as the next most influential observation). If we exclude this stimulus, we find a weaker correlation (*r =* 0.290; one-sided *t*-test; t(9) = 0.908, p = 0.194). Furthermore, there was no evidence of a relationship with ND when considering only the scrambled stimuli and their unscrambled counterparts (*r* = – 0.537; two-sided *t-*test; *t*(3) = –1.104. p *=* 0.350). Thus, we conclude that ND is not inherited from SD. We also did not find a relationship with stimulus luminance, contrast, or spectral energy (**Supplementary Figure 4**).

## 3 Discussion

Our results show that excitatory L2/3 neurons in higher visual areas AL and AM have more differentiated responses to movie stimuli with naturalistic structure than to phase-scrambled stimuli with closely matched low-order statistics, indicating that these populations are uniquely sensitive to high-level natural features in this stimulus set. We found this difference in neurophysiological differentiation (ND) at the level of single experimental sessions, and it was robust to complementary methods of measuring ND. Moreover, we found that effect sizes were larger with increasing pupil diameter and locomotion, suggesting that the increased ND in L2/3 of AL and AM is dependent on the animal’s arousal level and behavioral state. Decoding analysis showed a marked lack of area and layer specificity: stimulus category could be accurately decoded from the activity of most cell populations we surveyed. In addition to the differences between unscrambled and scrambled stimuli, we found differences in ND among unscrambled continuous stimuli. Finally, we argued that ND is not merely inherited from the differentiation of the stimuli.

The precise functional specialization of individual higher visual areas in the mouse, as well as that of V1, remains unclear (Glickfeld & Olsen, 2017). Recent large-scale anatomical (J. A. Harris et al., 2019) and functional (Siegle et al., 2019) studies of feedforward and feedback connectivity in the mouse visual system have uncovered a “shallow hierarchy” in which V1 lies at the base, followed by LM, RL, AL, and PM, with AM at the top. In this light, our findings that ND in L2/3 of AL and AM is sensitive to high-level naturalistic structure could be interpreted as a reflection of hierarchical processing, which may be constructing a richer dynamical repertoire for perception of naturalistic stimuli at higher hierarchical levels. Interestingly, we did not find this effect in PM, despite its intermediate position between AL and AM in the hierarchy, suggesting that such hypothetical processing towards richer repertoires is not fully determined by the one-dimensional hierarchy, but may involve specific pathways through subsets of higher visual areas. These observations indicate that differentiation analysis may be used to refine our understanding of functional specialization of these areas and uncover differences between them that can be used to direct further investigations and generate hypotheses.

A recent study found that feedback projections from higher visual areas to L2/3 excitatory neurons in V1 create a second receptive field (RF) surrounding the feedforward RF and that these RFs are mutually antagonistic, pointing to a role for these neurons in predictive processing (Keller et al., 2020). If this pattern is found at higher levels of the visual hierarchy, then the layer specificity of our findings could be explained by a scenario in which top-down feedback inputs to AL and AM from areas higher in the putative dorsal stream (Marshel et al., 2011; Wang et al., 2012) are integrated with feedforward inputs in L2/3 to compute and relay prediction errors about high-level visual features. In this scenario, the naturalistic stimuli, which contain high-level features that are presumably less predictable, would elicit more prediction errors and thus more differentiated activity.

Stimulus-evoked activity in cortex is powerfully modulated by arousal level and behavioral state (McGinley, Vinck, et al., 2015; Salkoff et al., 2020). Locomotion is associated with heightened arousal, increased membrane depolarization, increased firing rates, increased signal-to-noise ratio, and enhanced stimulus encoding (Bennett et al., 2013; Dadarlat & Stryker, 2017; Niell & Stryker, 2010; Polack et al., 2013; Vinck et al., 2015). Pupil diameter can serve as an index of arousal (Larsen & Waters, 2018), and exhibits an inverted-U relationship with task performance such that performance is optimal at intermediate arousal levels (McGinley, David, et al., 2015; McGinley, Vinck, et al., 2015). Larger pupils are associated with increases in the gain, amplitude, signal-to-noise ratio, and reliability of responses in V1 (Reimer et al., 2014). Thus, our finding that increased pupil diameter and locomotion activity are associated with larger effect sizes could be explained by an increase in response gain or amplitude in V1 that is inherited by downstream AL and AM: since the ND in these areas is selective for naturalistic structure, increased bottom-up drive could accentuate unscrambled-scrambled differences in ND.

Alternatively, response gain or amplitude in higher visual areas could be modulated directly by subcortical arousal systems. The noradrenergic and cholinergic systems are likely candidates, although it is not clear why noradrenergic modulation would cause an effect specific to L2/3; as for cholinergic modulation, Pafundo et al. (2016) showed that V1 and LM are differentially modulated by basal forebrain stimulation such that the response gain and reliability of excitatory L2/3 neurons was enhanced in V1 but not in LM, despite an even distribution of basal forebrain axon fibers across all layers in both areas. However, neuromodulatory regulation of activity in other higher visual areas, in particular AL and AM, has not yet been characterized in great detail and would be a fruitful topic for future studies. Another possibility is a top-down effect, in which increases in arousal and locomotion reflect increased cognitive or attentional engagement with the stimuli that favors processing of high-level stimulus features, selectively increasing ND for the unscrambled stimuli. In the passive viewing paradigm employed here, in which the animal is not motivated to attend to the stimuli, it is likely that top-down modulation of sensory processing varies considerably across the experimental session as arousal and attention fluctuate.

Though differentiation analysis revealed area- and layer-specific differences in responses to unscrambled and phase-scrambled stimuli, our ability to decode stimulus category from neural responses was remarkably similar across areas and layers. This contrast in our results highlights an important methodological distinction: decoding is a powerful means to reveal information content, but this information is necessarily measured from the extrinsic perspective (Buzsáki, 2019, Tononi, 2004; Oizumi et al., 2014; Tononi et al., 2016). The presence of information about a stimulus in a neural circuit does not imply that the information is functionally relevant to the system in question (Brette, 2019). As an extreme example, stimulus category would presumably be perfectly decodable from photons impinging on the retina, but this would reveal nothing of interest about perception. By contrast, ND is an intrinsic measure in the sense that it is defined without reference to a stimulus (Boly et al., 2015; Mensen et al., 2017, 2018). In the brain, a complex evolved system in which activity is energetically costly, ND may be a signature of functionally relevant dynamics. The dissociation we find between ND and decoding performance indicates that differentiation analysis can point to populations of interest that are not revealed by detecting stimulus-relevant information.

Finally, we also found that the predator stimulus and the “mousecam” stimulus elicited significantly higher ND than other unscrambled continuous stimuli. The predator stimulus finding is intriguing because that stimulus has lower luminance, contrast, and spectral energy than the clip of conspecifics in a home cage (**Supplementary Figure 4**). Given the importance of detecting natural predators, it is plausible that the high ND evoked by this stimulus reflects its particular salience to the visual system, driven by high-level features such as the presence of the predator rather than low-order stimulus statistics. In any case, this demonstrates that differentiation analysis can be used to probe differences in visual responses at the level of individual stimuli.

It is important to keep in mind the limitations of the data we collected. Firstly, calcium imaging provides only an imperfect proxy of neuronal activity: simultaneous juxtacellular electrophysiology indicates that the fluorescence signal from Ca^2+^ indicators is more sensitive to bursts of action potentials than sparse, low-frequency spiking (Chen et al., 2013; Huang et al., 2020; Ledochowitsch et al., 2019; Siegle et al., 2020; Wei et al., 2020). Such activity may contribute to ND but would not be present in this dataset. However, given the typically sparse response properties of L2/3 excitatory neurons compared to those in deeper layers (Barth & Poulet, 2012), it is possible that this limitation may only obscure even stronger L2/3 specificity. Secondly, for this exploratory study we opted to use a range of naturalistic stimuli and a limited number of phase-scrambled control stimuli in order to include diverse high-level features. Future studies measuring stimulus-evoked ND could test our findings using a larger set of artificial stimuli that control for other low-level stimulus characteristics, *e.g.* optical flow, in addition to the power spectrum. Thirdly, there was considerable variability in arousal state and locomotor activity in our passive viewing paradigm. Given these factors’ modulation of effect size, future work might uncover larger effects by employing an active paradigm in which the animal is motivated by reward to attend to the stimuli.

In summary, we measured stimulus-evoked differentiation of neural activity with cellular resolution and found increased ND in response to unscrambled versus scrambled stimuli. This effect was driven by L2/3 excitatory cells in AL and AM and was enhanced at higher arousal levels. To our knowledge, the present study is the first to systematically measure stimulus-evoked differentiation with cellular resolution across multiple cortical areas and layers. These results advance our understanding of the functional differences among higher visual areas, and future work should seek to integrate our findings into the emerging picture of a shallow hierarchy in the mouse visual system, for example by investigating potential differences in neuromodulation among these areas or the contrast between AL/AM and PM. Differentiation analysis is motivated by IIT, and provides an “intrinsic” analytical approach that can complement “extrinsic” measures such as decoding performance, which in this dataset did not distinguish specific cell populations. This method can be used to compare individual stimuli and may provide a readout of the degree to which a given stimulus induces a rich and varied perceptual experience. Future studies should investigate stimulus-evoked differentiation with cellular resolution in humans (and perhaps non-human primates), where subjective reports are available, and thereby determine the relative contributions of distinct cell populations to ND while correlating ND with phenomenology.

## 4 Methods

The AIBS optical physiology pipeline is described in detail in de Vries et al. (2020) and Groblewski et al. (2020). Analysis was performed with custom Python and R code using `numpy` (C. R. Harris et al., 2020), `scipy` (Virtanen et al., 2020), `pandas` (Reback et al., 2020), `scikit-learn` (Pedregosa et al., 2011), `matplotlib` (Hunter, 2007), `seaborn` (Waskom & the seaborn development team, 2020), `lme4` (Bates et al., 2015, p. 4), `multcomp` (Hothorn et al., 2008), and `emmeans` (Lenth, 2020).

### 4.1 Transgenic mice

All animal procedures were approved by the Institutional Animal Care and Use Committee at the AIBS. We maintained all mice on reverse 12-hour light cycle following surgery and throughout the duration of the experiment and performed all experiments during the dark cycle. We used the transgenic mouse line Ai93, in which GCaMP6f expression is dependent on the activity of both Cre recombinase and the tetracycline controlled transactivator protein (tTA) (Madisen et al., 2010). Triple transgenic mice (Ai93, tTA, Cre) were generated by first crossing Ai93 mice with Camk2a-tTA mice, which preferentially express tTA in forebrain excitatory neurons.

Cux2-CreERT2;Camk2a-tTA;Ai93(TITL-GCaMP6f) expression is regulated by the tamoxifen-inducible Cux2 promoter, induction of which results in Cre-mediated expression of GCaMP6f predominantly in superficial cortical layers 2, 3 and 4. Rorb-IRES2-Cre;Cam2a-tTA;Ai93 exhibit GCaMP6f in excitatory neurons in cortical layer 4 (dense patches) and layers 5 & 6 (sparse). Rbp4-Cre;Camk2a-tTA;Ai93 exhibit GCaMP6f in excitatory neurons in cortical layer 5.

### 4.2 Surgery

Transgenic mice expressing GCaMP6f were weaned and genotyped at ~P21, and surgery was performed between P37 and P63. The craniotomy was centered at X = –2.8 mm and Y = 1.3 mm with respect to lambda (centered over the left mouse visual cortex). A circular piece of skull 5 mm in diameter was removed, and a durotomy was performed. A coverslip stack (two 5 mm and one 7 mm glass coverslip adhered together) was cemented in place with Vetbond. Metabond cement was applied around the cranial window inside the well to secure the glass window.

### 4.3 Intrinsic imaging

A retinotopic map was created using intrinsic signal imaging (ISI) in order to define visual area boundaries and target in vivo two-photon calcium imaging experiments to consistent retinotopic locations. These maps were generated while mice were lightly anesthetized with 1–1.4% isoflurane. See de Vries et al. (2020) for a complete description of this procedure and related processing steps.

### 4.4 Habituation

Following successful ISI mapping, mice spent two weeks being habituated to head fixation and visual stimulation. During the second week, mice were head-fixed and presented with visual stimuli, starting for 10 minutes and progressing to 50 minutes of visual stimuli by the end of the week. During this week they were exposed to the “mouse montage 2” stimulus (see **Stimuli**).

### 4.5 Imaging

Calcium imaging was performed using a two-photon-imaging instrument (Nikon A1R MP+). Laser excitation was provided by a Ti:Sapphire laser (Chameleon Vision – Coherent) at 910 nm. Mice were head-fixed on top of a rotating disc and free to run at will. The screen center was positioned 118.6 mm lateral, 86.2 mm anterior and 31.6 mm dorsal to the right eye. The distance between the screen and the eye was 15 cm. Movies were recorded at 30 Hz using resonant scanners over a 400 μm field of view.

Excitatory neurons from cortical L2/3, L4, and L5 were imaged (L2/3: 3 mice, 15 sessions; L4: 3 mice, 15 sessions; L5: 3 mice, 14 sessions) in 5 visual areas: V1 (9 sessions), L (9 sessions), AL (8 sessions), AM (9 sessions), and PM (9 sessions).

### 4.6 Behavioral data

#### 4.6.1 Locomotion

Locomotion velocity data recorded from the running wheel were preprocessed as follows. First, artifacts were removed using custom code that iteratively identified large positive or negative peaks (indicative of artifactual discontinuities in the signal) in several passes of `scipy.signal.find_peaks` (specific parameters were manually chosen for each session). Remaining artifacts were then manually removed by inspecting the resulting timeseries and visually identifying clear discontinuities. The removed samples were filled using linear interpolation (`pandas.Series.interpolate`).

The resulting signal was then low-pass filtered at 1 Hz using a zero-phase 4^th^-order Butterworth filter (`scipy.signal.butter(2, 1/15, btype=’lowpass’, output=’ba’, analog=False`) applied with `scipy.signal.filtfilt`).

For the analysis in Figure 4, the fraction of time spent running was computed by binarizing the preprocessed velocity timeseries at a threshold of 2.5 cm/s.

#### 4.6.2 Pupillometry

Pupil diameter was extracted from video of the mouse’s ipsilateral eye (relative to the stimulus presentation monitor) using the AllenSDK (https://github.com/AllenInstitute/AllenSDK) as described in de Vries et al. (2020).

Briefly, for each frame of the video an ellipse was fitted to the region corresponding to the pupil as follows: a seed point within the pupil was identified via convolution with a black square; 18 rays were drawn starting at this seed point, spaced 20 degrees apart; the candidate boundary point between the pupil and iris along that ray was identified by a change in pixel intensity above a session-specific threshold; a RANSAC algorithm was used to fit the an ellipse to the candidate boundary points using linear regression with a conic section constraint; and fitted parameters of the regression were converted to ellipse parameters (coordinates of the center, lengths of the semi-major and semi-minor axes, and angle of rotation with respect to the x-axis). Pupil diameter was taken to be twice the semi-major axis of the fitted ellipse.

The resulting timeseries contained some artifacts, which we removed by the same combination of automated and manual methods used for the locomotion timeseries (see **Locomotion**).

For the analysis shown in Figure 4, each pupil diameter timeseries was normalized by dividing by the maximum diameter that occurred during stimulus presentations.

### 4.7 Stimuli

We created twelve 30 s greyscale naturalistic and artificial movie stimuli.

The eight naturalistic stimuli (**Figure 9**, top) consisted of three montages of six 5 s clips, spliced together with jump cuts, and four continuous stimuli. The “mouse montage 1” stimulus contained clips of conspecifics, a snake, movement at ground level through the underbrush of a wooded environment, and a cat approaching the camera. The “mouse montage 2” stimulus contained different footage of movement through the wooded environment; different footage of a cat approaching the camera; conspecifics in a home cage filmed from within the cage; crickets in a home cage filmed from within the cage; footage of the interior of the home cage with environmental enrichment (a shelter, running wheel, and nesting material); and a snake filmed at close range orienting towards the camera. The “human montage” contained clips of a man talking animatedly to an off-screen interviewer; a café table where food is being served; automobile traffic on a road viewed from above; a woman in the foreground taking a photo of a city skyline; footage of a road filmed from the passenger seat of a vehicle; and a close shot of a bowl of fruit being tossed. The four continuous stimuli were: footage of a snake at close range orienting towards the camera; crickets in a home cage filmed from within the cage; a man writing at a table; movement through a wooded environment at ground level; and conspecifics in a home cage. No two stimuli contained identical clips.

**Figure 9.**
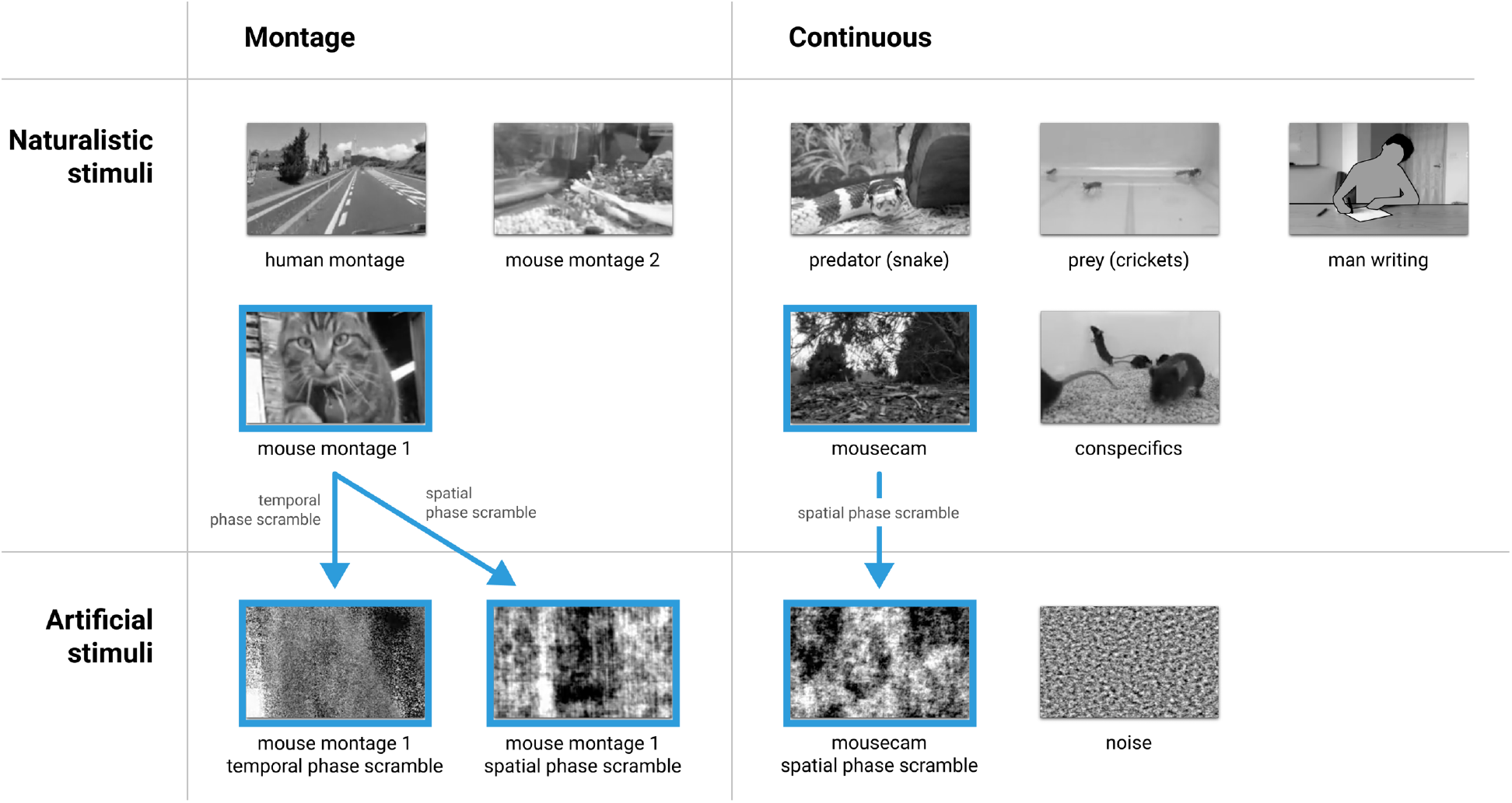
Stimuli. Twelve 30 s long greyscale naturalistic (top) and artificial (bottom) movie stimuli were presented. Left: montages of six 5 s clips; right: continuous 30 s clips. Stimuli used in the main analysis are outlined in blue. Arrows indicate the phase-scrambling procedures. *Note:* the “man writing” stimulus frame has been de-identified for presentation in this preprint in accordance with bioRxiv policy.

The four artificial stimuli (**Figure 9**, bottom) consisted of two phase-scrambled versions of the “mouse montage 1” stimulus, a phase-scrambled version of the “mousecam” stimulus (see **Phase scrambling**), and a high-pass-filtered 1/*f* noise stimulus.

The stimuli were presented in a randomized block design with 10 repetitions, with 4 s of static mean-luminance grey presented between stimuli (**Figure 1F**). 60 s of mean-luminance grey (to record spontaneous activity) and a 60 s high-contrast sparse noise stimulus were also presented in the beginning of each session (this stimulus was not analyzed in this work).

#### 4.7.1 Phase scrambling

Two methods of phase scrambling were used: temporal and spatial, described in detail below. Briefly, for the temporal scrambling we independently randomized the phase of each pixel’s intensity timeseries in contiguous, nonoverlapping windows of 1 s. For the spatial scrambling, we randomized the phase of the spatial dimensions of the three-dimensional spectrum of each window. The “mouse montage 1” stimulus was phase-scrambled using both procedures to obtain the “mouse montage 1, temporal phase scramble” and “mouse montage 1, spatial phase scramble” stimuli. The “mousecam” stimulus was scrambled using the spatial procedure to obtain the “mousecam, spatial phase scramble” stimulus.

##### Temporal phase scramble

First, the stimuli were windowed into contiguous, nonoverlapping 1 s segments (30 frames each). For each 1 s window, we applied the following procedure:

We estimated the one-dimensional spectrum of each pixel’s intensity timeseries with the discrete Fourier transform (DFT) using the NumPy function `numpy.fft.fft`. The phase and magnitude of each spectrum were computed with `numpy.angle` and `numpy.abs` respectively. For each pixel, we generated a 14-element random vector drawn uniformly from the interval [0, 2π]. A randomized phase was then obtained for that pixel by concatenating the first element of the original phase, the random vector, the 15^th^ element of the original phase, and the negative reversed random vector. This yielded a 30-element phase vector with the required conjugate symmetry of the spectrum of a 1 s real-valued signal sampled at 30 frames per second. The randomized phase was then combined with the spectral magnitude and transformed back into the time domain with the inverse DFT using `numpy.fft.ifft`, yielding a temporally phase-scrambled version of that pixel’s intensity timeseries. Each pixel’s timeseries was independently phase-scrambled in this fashion.

This resulted in 30 independently phase-scrambled 1 s windows. These windows were then concatenated to obtain the full 30 s temporally phase-scrambled stimulus.

##### Spatial phase scramble

First, the stimuli were windowed into contiguous, nonoverlapping 1 s segments (30 frames each). For each window, we applied the following procedure. The three-dimensional Fourier spectrum (frame, width, and height) was estimated with the DFT using `numpy.fft.fftn`. The phase and magnitude of the spectrum were computed with `numpy.angle and numpy.abs` respectively. To randomize the phase in the spatial dimensions, we generated a random signal in the time domain with the same dimensions as a stimulus frame (192 pixels wide by 120 pixels high) and computed its phase in the frequency domain as described above. This two-dimensional random spatial phase was added to the spatial dimensions of the three-dimensional stimulus phase. After being randomized in this way, the stimulus phase was recombined with the spectral magnitude and transformed back into a time-domain signal with the inverse DFT using `numpy.fft.ifftn`. The 30 resulting phase-scrambled 1 s windows were then concatenated to obtain the full 30 s spatially phase-scrambled stimulus.

##### Effect of phase-scrambling

The greyscale movie stimuli were represented in the stimulus presentation software as arrays of unsigned 8-bit integers. The limitations of this representation resulted in phase-scrambled stimuli with power spectra that were close but not identical to the power spectrum of their unscrambled counterparts.

Specifically, although the phase scrambling procedures described above leave the power spectrum unchanged, they do not necessarily preserve the range of the resulting real-valued signal. In our case, applying these procedures to our stimuli resulted in phase-scrambled stimuli in which the pixel intensities occasionally lay outside the range [0, 255]. Thus, in order to represent the phase-scrambled stimuli with 8-bit integers, we truncated the result so that negative intensities were set to 0 and intensities greater than 255 were set to 255. This operation does affect the power spectra, and as a result the spectra of the unscrambled and scrambled stimuli are closely matched but not equal.

### 4.8 Differentiation analysis

#### 4.8.1 Spectral differentiation

Our analysis of the responses to the stimuli follows the techniques developed in previous work in humans (Boly et al., 2015; Mensen et al., 2017, 2018). The spectral differentiation measure of ND used by Mensen et al. (2018) was designed for analysis of timeseries responses to continuous movie stimuli, and was found to be positively correlated with subjective reports of stimulus “meaningfulness”. We employed this measure with our Ca^2+^ imaging data: (A) for each cell, the ΔF/F_0_ trace of each cell during stimulus presentation was divided into 1 s windows; (B) the power spectrum of each window was estimated using a Fourier transform; (C) the “neurophysiological state” during each 1 s window was defined as a vector in the high-dimensional space of cells and frequencies (*i.e.*, the concatenation of the power spectra in that window for each cell); (D) the ND in response to a given stimulus was calculated as the median of the pairwise Euclidean distances between every state that occurred during the stimulus presentation. A schematic illustration is shown in **Figure 2**.

We normalized spectral differentiation values by the square root of the number of cells in the recorded population, reasoning as follows. Consider a hypothetical population of cells that each exhibit the same temporal pattern of activity. The spectral differentiation of such a population will be proportional to the square root of its size, because the Euclidean distance is used to compare neurophysiological states. If we have two such populations differing only in the number of cells, their activity should be considered to be equally differentiated for our purposes, since their temporal patterns are identical; any differences in spectral differentiation would be due to the (arbitrary) number of cells captured in the imaging session. Thus, we divided by the square root of the population size to remove this dependency.

#### 4.8.2 Multivariate differentiation

We also measured ND using a multivariate approach that considers spatiotemporal differences in activity patterns. For each experimental session, we extracted ΔF/F_0_ traces recorded during all stimulus presentations and concatenated them to obtain an *m* × *n* matrix of responses where *m* is the number of two-photon imaging samples and *n* is the number of traces. This matrix was then downsampled by summing the ΔF/F_0_ traces within 100 ms bins. We used a nonlinear dimensionality reduction procedure, Uniform Manifold Approximation and Projection for Dimension Reduction (Python package `umap-learn`, McInnes et al., 2018), to reduce this matrix to *m* × 10 with parameters `UMAP(n_components=10, metric=“euclidean”, n_neighbors=100, min_dist=0.0)`. Each row of the resulting matrix was a 10-dimensional vector that represented the state of the cell population during the corresponding 100 ms interval. We then grouped the rows of the resulting matrix by stimulus. Each row vector can be thought of as a point in ℝ^10^, so that each stimulus was associated with a cloud of points corresponding to the population states that the stimulus evoked over the course of all 10 trials.

The intuition motivating this approach is that we can operationalize the notion of neurophysiological differentiation by measuring the dispersion of this point cloud. The more distant two points are, the more different are the corresponding responses of the cell population; thus, if a stimulus evokes many different population states, the point cloud will be more spread out in response space. Therefore, we measured ND evoked by each stimulus by finding the centroid of its associated point cloud and taking the mean Euclidean distance of each point to the centroid.

### 4.9 Statistics

#### 4.9.1 Linear mixed effects models

For aggregate statistics across all experimental sessions, we employed linear mixed effects models using the `lmer` function from the `lme4` package in R with `REML = FALSE` (Bates et al., 2015, p. 4). The distributions of ND values for both spectral and multivariate differentiation measures were well-approximated by log-normal distributions, so we applied a logarithmic transformation to ND values prior to statistical modeling.

First we fit an LME model with cortical layer, stimulus category (unscrambled or scrambled), and their interaction as fixed effects, with experimental session as a random effect (`lme4` formula: “`differentiation ~ 1 + layer * stimulus_category + (1 | session)`”). To test-layer specificity, we then fit a reduced model with the interaction removed (“`differentiation ~ 1 + area + stimulus_category + (1 | session)`”) and used a likelihood ratio test to compare the two models.

Next we fit an LME model with cortical area, stimulus category, and their interaction as fixed effects, with experimental session as a random effect (`lme4` formula: “`differentiation ~ 1 + area * stimulus_category + (1 | session)`”). To test-area specificity, we fit a reduced model with the interaction removed (“`differentiation ~ 1 + area + stimulus_category + (1 | session)`”) and used a likelihood ratio test to compare the two models.

We tested for differences among the unscrambled continuous stimuli (“snake (predator)”, “crickets (prey)”, “man writing”, “mousecam”, and “conspecifics”) by fitting an LME model with stimulus as a fixed effect and experimental session as a random effect (`lme4` formula: “`differentiation ~ 1 + stimulus + (1 | session)`”).

#### 4.9.2 Post hoc tests

Post hoc one-sided *z*-tests of layer and area specificity were performed calling the `glht` function from the `multcomp` package in R on each LME model with contrasts between stimulus categories (unscrambled or scrambled) within each layer and area, respectively. P values were adjusted for multiple comparisons using the single-step method in `multcomp` (Hothorn et al., 2008).

Post hoc two-sided *t*-tests for pairwise differences among the unscrambled continuous stimuli were performed with the `emmeans` function from the `emmeans` package in R (“`emmeans(model, pairwise ~ stimulus`”), with p values adjusted for multiple comparisons using Tukey’s method (Lenth, 2020).

#### 4.9.3 Permutation tests

Permutation tests were performed for each experimental session to test whether spectral differentiation evoked by unscrambled stimuli was greater than that evoked by scrambled stimuli. We obtained a null distribution by randomly permuting the trial labels (unscrambled or scrambled) 20,000 times and computing the difference in mean spectral differentiation on unscrambled and scrambled trials for each permutation. P values were computed as the fraction of permutations for which the permuted difference was greater than the observed difference, and significance is reported at the level of α = 0.05.

### 4.10 Decoding analysis

For each experimental session, we decoded stimulus category (unscrambled or scrambled) using linear discriminant analysis with the Python package `scikit-learn` (Pedregosa et al., 2011). First, the responses to each category were concatenated to form a *s* × (*n · t*) matrix, where *s* is the number of stimulus presentation trials, *n* is the number of cells recorded, and *t* is the number of two-photon imaging samples in a single trial. To obtain a tractable number of features for linear discriminant analysis, we used PCA to reduce the dimensionality of the matrix such that the number of components *c* was sufficient to retain 99% of the variance along the rows, yielding an *s* × *c* matrix (`sklearn.decomposition.PCA(n_components=0.99)`). This was then used to train a shrinkage-regularized LDA classifier with fivefold cross-validation (`sklearn.discriminant_analysis.LinearDiscriminantAnalysis(solver=’lsqr’, shrinkage=’auto’)`). We report the mean balanced accuracy score (`sklearn.metrics.balanced_accuracy_score`) on the heldout test data across cross-validation folds. Chance performance is 0.5.

For the analysis shown in Supplementary Figure 2, we used the same procedure as described above, but the classifier was trained to decode stimulus identity rather than category; chance performance is 1/12. For Supplementary Figure 3, we used the same procedure but trained the classifier using only responses to the 5 continuous naturalistic stimuli, and classifier performance was evaluated for each stimulus separately with the F1 score.

## Acknowledgements

We thank Andrew Haun for helpful comments and assistance with creating the noise stimulus; Jonathan Lang for assistance filming the naturalistic stimuli; and Larissa Albantakis, Leonardo S. Barbosa, Tom Bugnon, Graham Findlay, Marcello Massimini, and Shuntaro Sasai for helpful discussions.

W.M. received support from the Natural Sciences and Engineering Research Council of Canada (RGPIN-2019-05418).

The data presented herein were obtained at the Allen Brain Observatory as part of the OpenScope project, which is operated by the Allen Institute. This work was supported by the Allen Institute, the Tiny Blue Dot Foundation, and in part by the Falconwood Foundation. We thank Allan Jones for providing the crucial environment that enabled our large-scale team effort. We thank the Allen Institute founder, Paul G. Allen, for his vision, encouragement, and support.

## Author Contributions

**William G. P. Mayner** Conceptualization, Data Curation, Formal Analysis, Investigation, Methodology, Software, Validation, Visualization, Writing – Original Draft Preparation, Writing – Review & Editing

**William Marshall** Conceptualization, Formal Analysis, Investigation, Methodology, Supervision, Validation, Writing – Review & Editing

**Yazan N. Billeh** Data Curation, Investigation, Methodology

**Saurabh R. Gandhi** Data Curation, Formal Analysis, Investigation, Validation, Visualization, Writing – Review & Editing

**Shiella Caldejon** Data Curation, Investigation, Project Administration

**Andrew Cho** Investigation

**Fiona Griffin** Data Curation, Investigation

**Nicole Hancock** Investigation

**Sophie Lambert** Investigation

**Eric Lee** Data Curation, Investigation, Software

**Jennifer Luviano** Investigation, Project Administration

**Kyla Mace** Investigation

**Chelsea Nayan** Investigation

**Thuyanh Nguyan** Data Curation, Investigation

**Kat North** Investigation

**Sam Seid** Data Curation, Investigation

**Ali Williford** Investigation

**Chiara Cirelli** Supervision, Writing – Review & Editing

**Peter Groblewski** Project Administration, Supervision

**Jerome Lecoq** Data Curation, Funding Acquisition, Methodology, Project Administration, Resources, Software, Supervision, Writing – Review & Editing

**Giulio Tononi** Conceptualization, Funding Acquisition, Resources, Supervision, Writing – Review & Editing

**Christof Koch** Conceptualization, Funding Acquisition, Resources, Supervision, Writing – Review & Editing

**Anton Arkhipov** Conceptualization, Funding Acquisition, Methodology, Project Administration, Resources, Supervision, Validation, Writing – Review & Editing

All authors reviewed and approved the final version of the manuscript.

## Supplementary figures

**Supplementary Figure 1.**
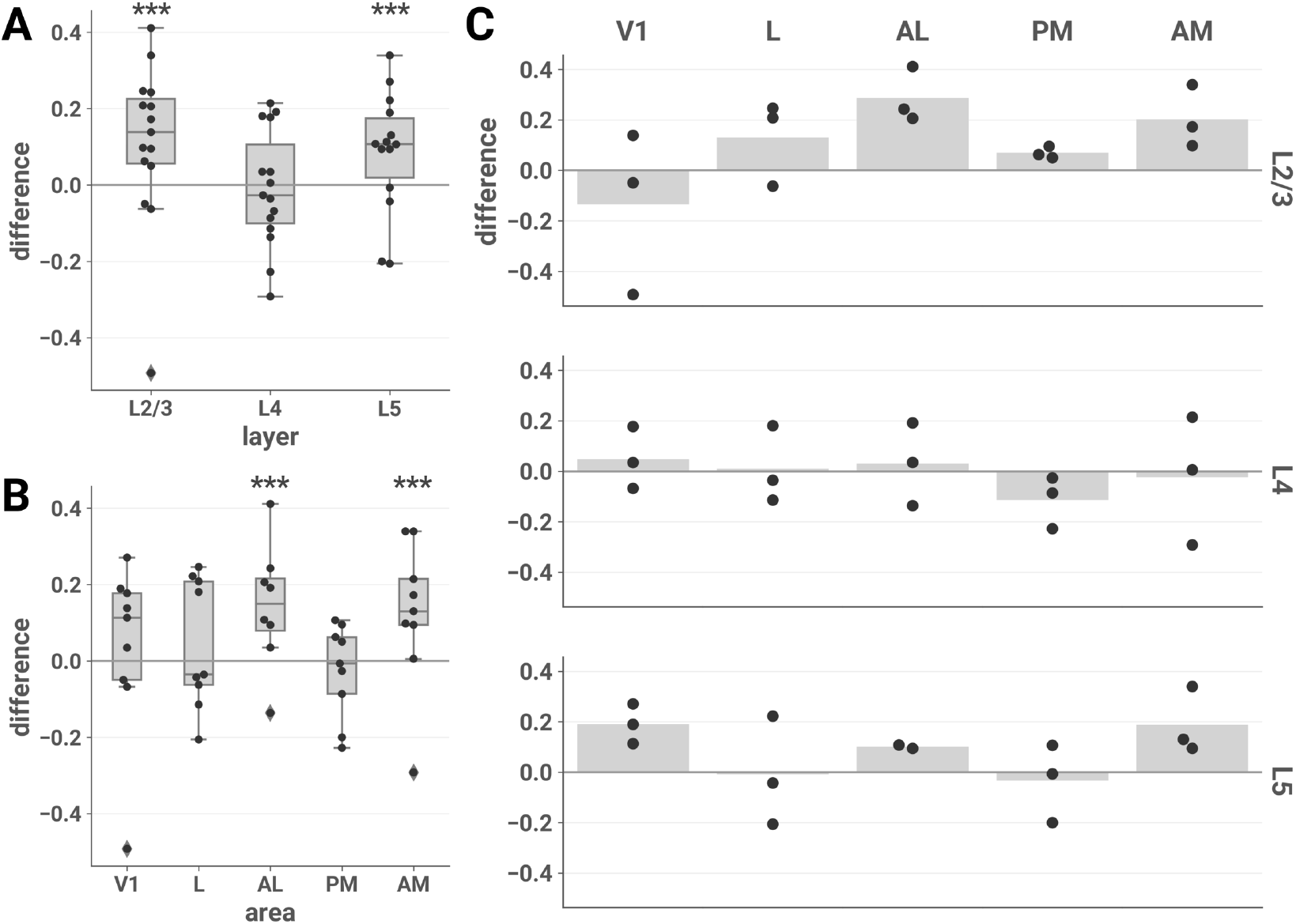
Naturalistic vs. artificial differences in ND across the entire stimulus set. The mean difference in ND of responses to all 8 naturalistic vs. all 4 artificial stimuli is plotted for each session by layer (A), area (B), and layer-area pair (C). Results are similar to the unscrambled vs. scrambled contrast shown in **Figure 2**. In this analysis, post hoc tests showed a significant effect also in L5; however, this contrast does not control for low-level stimulus characteristics and is thus harder to interpret. **(A)** We fit an LME model with stimulus category (naturalistic or artificial), layer, and their interaction as fixed effects and found a significant interaction (likelihood ratio test, *χ^2^*(2) = 16.343, p = 0.000283). Post hoc one-sided *z*-tests (adjusted for multiple comparisons): L2/3, z = 4.974, p = 9.82e–7; L4, z = –0.450, p = 0.965; L5, z = 3.745, p = 0.000271. **(B)** We fit an LME model with stimulus category (naturalistic or artificial), area, and their interaction as fixed effects and found a significant interaction (likelihood ratio test, *χ^2^*(2) = 16.343, p = 0.000283). Post hoc one-sided *z*-tests (adjusted for multiple comparisons): V1, z = 1.207, p = 0.725; L, z = 1.523, p = 0.495; AL, z = 4.715, p = 1.21e–5; PM, z = –0.907, p = 0.896; AM, z = 4.249, p = 0.000107). **(A)** and **(B)**: asterisks indicate significant post hoc tests in the layer (A) and area (B) interaction LME models (***, p < 0.001). Boxes indicate quartiles; whiskers indicate the minimum and maximum of data lying within 1.5 times the inter-quartile range of the 25% or 75% quartiles; diamonds indicate observations outside this range. **(C)** Mean values are indicated by bars.

**Supplementary Figure 2.**
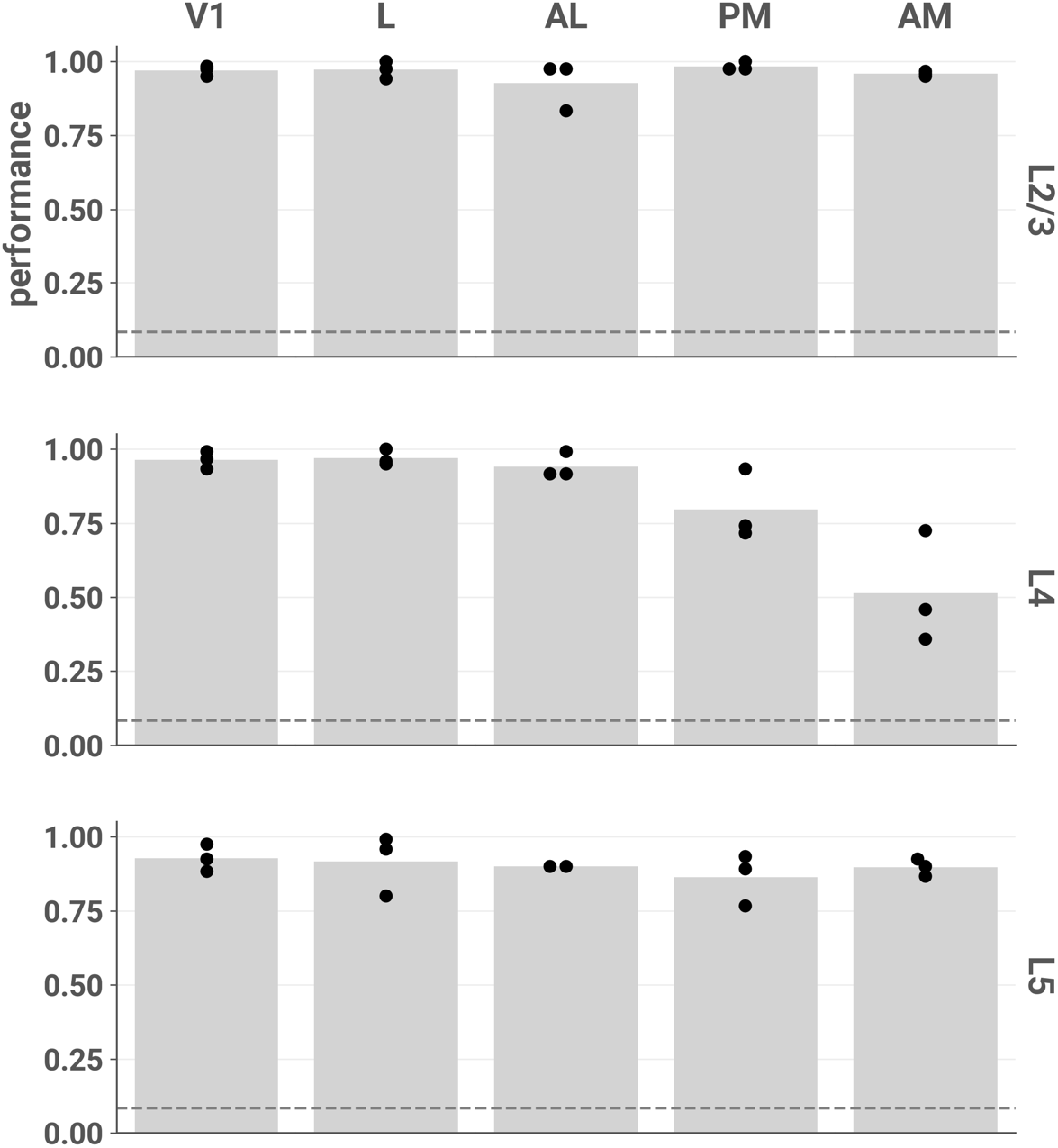
Stimulus identity can be accurately decoded from most layers and areas using responses to all 12 stimuli. Each point represents the mean fivefold cross-validated balanced accuracy score of linear discriminant analysis performed on a single session (see **Decoding analysis)**. Chance performance is 1/12, indicated by the dotted line.

**Supplementary Figure 3.**
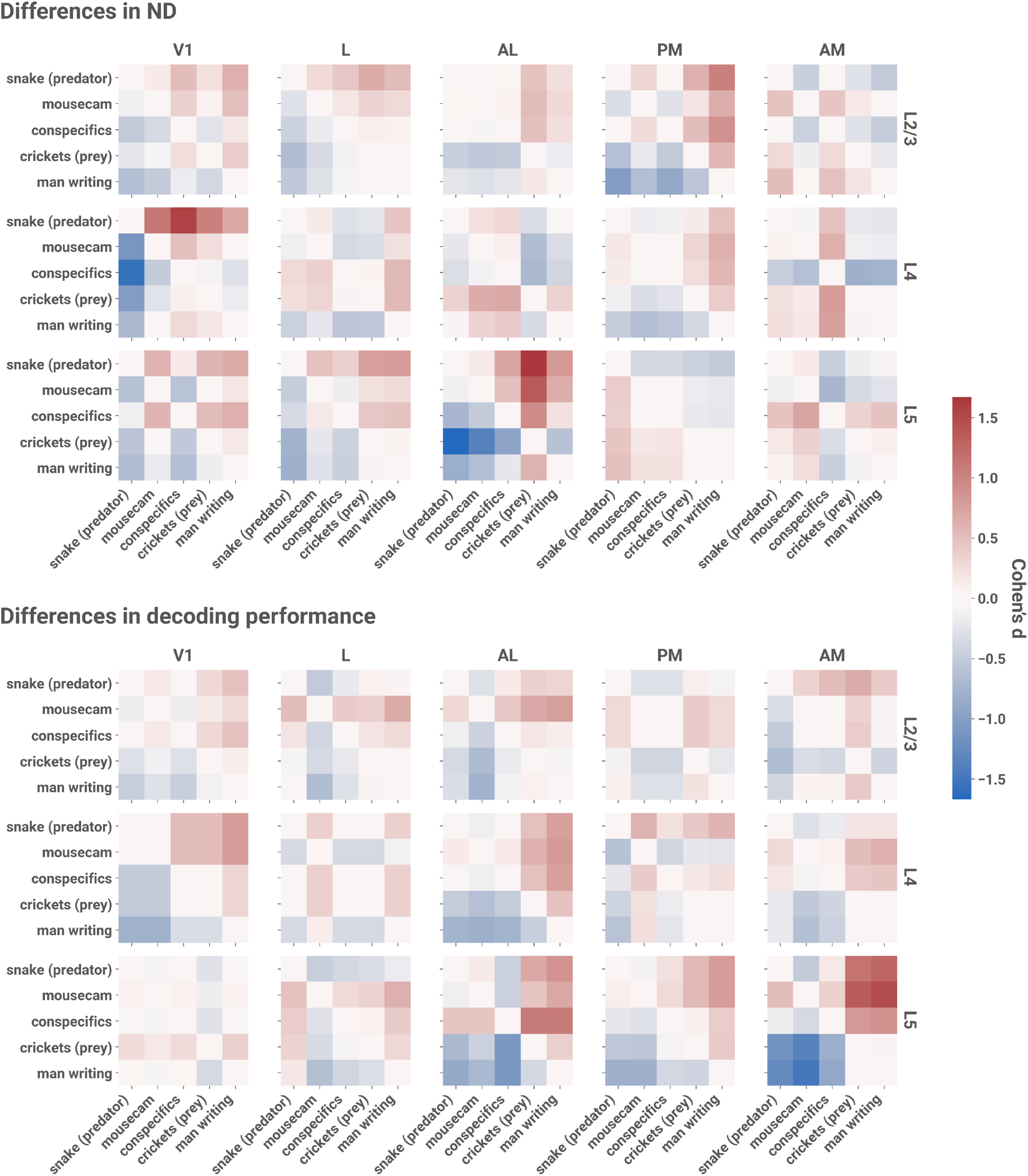
Within-category differences in ND vs. within-category differences in decoding performance, by layer and area. Top: Cohen’s *d* for pairwise mean differences in ND among naturalistic stimuli without jump cuts. Bottom: Cohen’s *d* for pairwise mean differences in stimulus identity decoding performance. For each session, we trained a linear discriminant analysis classifier using only responses to these 5 stimuli; classification performance was evaluated as the mean fivefold cross-validated F1 score for each stimulus (see **Decoding analysis**).

**Supplementary Figure 4.**
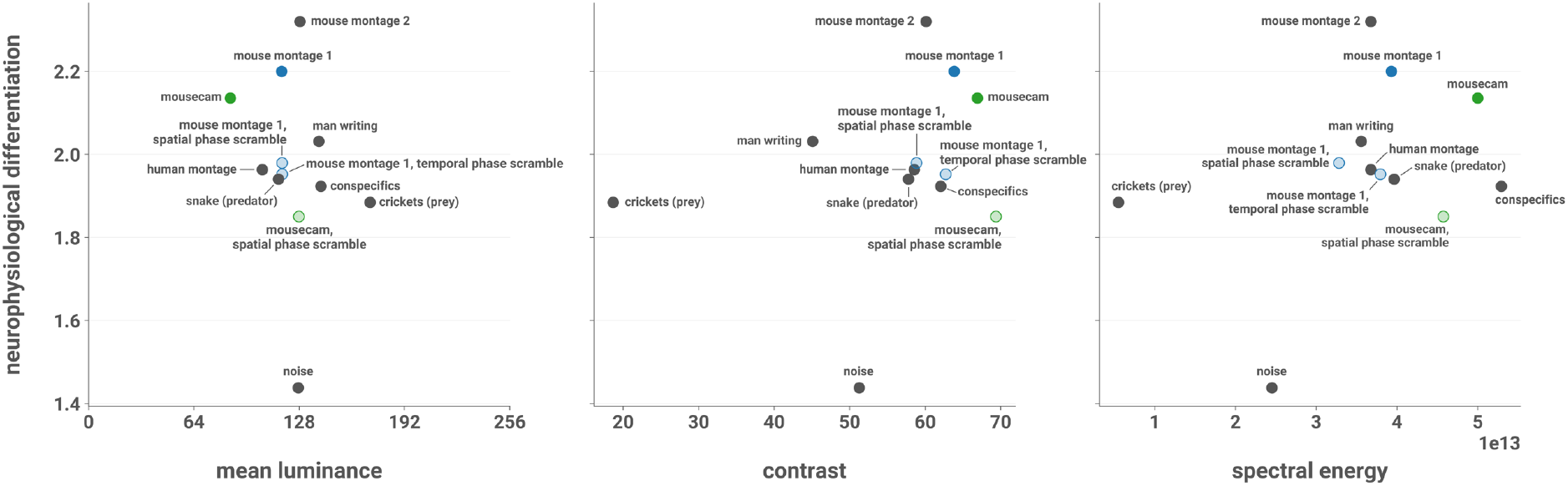
ND vs. low-level stimulus characteristics. ND is plotted against the mean luminance, contrast, and spectral energy of the stimuli. Mean luminance was computed as the average pixel intensity. Contrast was calculated as the standard deviation of pixel intensities. Spectral energy was computed as the sum of the energy spectral density of each pixel’s intensity timeseries after removing the DC component.

